# Patterns and mechanisms underlying ecoregion delineation in North American freshwater plants

**DOI:** 10.1101/2021.03.25.436944

**Authors:** Janne Alahuhta, Jorge García–Girón

## Abstract

**Aim:** Biogeographical regionalisations are actively studied in different ecosystems, because they increase our understanding on fundamental broad□scale patterns and can help us in the establishment of conservation areas. Thus, we studied how well existing freshwater ecoregions describe geographical delineation for inland water plants and which ecogeographical gradients explain them.

**Location:** North America, excluding Mexico and remote islands.

**Taxon:** Freshwater vascular plants of all taxa and different functional groups.

**Methods:** Using newly available fine–grained data on freshwater plant distributions, we calculated internal homogeneity and cross–boundary heterogeneity among neighbouring ecoregions. We further integrated measures of community dissimilarity to assess whether the degree of within–ecoregion homogeneity and distinctness are driven by their relationships to species replacements and richness differences, and explored how a complex suite of ecogeographical mechanisms and plant life forms affect ecoregion delineation using spatially explicit regression routines.

**Results:** We found a clear geographical patterning of ecoregion robustness for North American freshwater plants, with their communities being more internally homogeneous and more similar to one another in polar and subtropical inland waters. Surprisingly, the degree of internal homogeneity and ecoregion distinctness were almost equally driven by species replacements and richness differences. Considering different life forms, ecoregion delineation performed best for emergent and floating–leaved plants. Finally, within–ecoregion homogeneity and distinctness were best explained by annual mean temperature and terrain ruggedness, respectively, with mean water alkalinity, ecoregion area and Late Quaternary Ice Age legacies having supplementary effects.

**Main conclusions:** Our findings emphasise that geographical regionalisations founded on a particular organismal group are not applicable for all taxa. Our study is a promising starting point for further investigations of geographical delineations for different freshwater taxa. These updated regionalisations can then be used for conserving different biotas in freshwaters, which are currently among the most threatened ecosystems in the world.

**Statement of significance:** In biogeographical regionalisation biota is categorized to meaningful geographical units, such as ecoregions. However, ecoregions delineated for a particular group of organisms may not be applicable to another assemblages. We studied how ecoregions founded on fish are suitable for freshwater plants across North America. Our findings suggest that these ecoregions did not show consistent robustness for freshwater plants in North America. This study is a promising starting point for further investigations of geographical delineations for different freshwater taxa, having also value in conservation planning and management.

## Introduction

Biogeographical regionalisation, which refers to classification of biotas into meaningful geographical units, is still one of the central objectives in biogeography and ecology (Ficetola et al., 2017; Kreft & Jetz, 2010; Smith et al., 2018). In these biogeographical units, such as biomes and ecoregions, taxonomic composition ought to be maximally homogenised inside their boundaries (i.e., species composition across an entire region is relatively consistent), while showing highest differences among neighbouring units (i.e., communities existing in different ecoregions are relatively dissimilar) (Divisek et al., 2016; Holt et al., 2013; Bailey, 2004). Regionalisation does not only help us to understand fundamental biogeographical patterns, including community interactions and ecosystem functioning, but also offers vital applied perspectives related to establishment of science–based conservation plans designed to protect regions, habitat types and taxa (Ennen et al., 2020; Dinerstein et al., 2017; Divisek et al., 2016; Droissart et al., 2018). A common approach in conservation is to identify biodiversity hotspots and ecoregions, where protection and restoration efforts are deliberately focused. However, delineation of geographical units has mostly focused on terrestrial taxa (e.g., Ficetola et al., 2017; Holt et al., 2013; Dinerstein et al., 2017; Higgins et al., 2016), whereas information on freshwater ecoregions has received less attention (but see Abell et al., 2008; Matamoros et al., 2016; Ennen et al., 2020). Moreover, available freshwater ecoregion definitions are typically founded on well–investigated assemblages, such as fish (Abell et al., 2008; Matamoros et al., 2016) and amphibians (Dias–Loyola et al., 2008), but they may not mirror biogeographical units for all freshwater organisms, not least because diversity patterns and distributions are not often correlated among taxonomic groups (Rolls et al., 2017; Ennen et al., 2020).

Recent advances have questioned whether ecoregions can effectively conserve biodiversity across regions and taxa (MacDonald, 2005; Smith et al., 2018). Ability of ecoregions to capture all variability inside them is uncertain because biogeographical units are not intrinsic properties of the biosphere (MacArthur, 1972). Distinction of ecoregions is also often founded on imprecise evaluation or qualitative expert judgements (Ennen et al., 2020). Furthermore, strength of ecoregion boundaries may vary among regions, organismal groups and taxonomic resolutions (Ficetola et al., 2017; Ennen et al., 2020; Smith et al., 2020). For example, Smith et al. (2020) found that ecoregions based on plants, animals and fungi were more distinct in tropical zones compared with other land areas. Similarly, ecoregion delineations also became less evident when the number of fish families was increased in the continental United States (Matamoros et al., 2016). These potential challenges in biogeographical regionalisation clearly imply that existing ecoregion distinctness founded on limited biota does not necessarily portray biogeographical units for all organismal groups and realms. As ecoregion approach can provide a useful tool in defining ecological and phytosociological communities (Blasi & Frondoni, 2011), it is highly important to assess whether current ecoregion classification schemes are valid for different biotas and ecosystem types, and which mechanisms modulate the robustness of neighbouring boundaries for these organisms.

Here, we present an analysis of the descriptive power of freshwater ecoregions across North America (25°N–78°N) at a 50 x 50 km spatial resolution founded on understudied freshwater plants. Capitalising on the troves of newly available fine–grained data on freshwater plant distributions (Alahuhta et al., 2020, Vieira et al., 2021) and ecoregion maps (Abell et al., 2008; FEOW), we integrate measures of community dissimilarity (Carvalho et al., 2012) with data on putative ecogeographical mechanisms and plant life forms (Sculthorpe, 1967; Cook, 1999) potentially underlying ecoregion delineation. More specifically, we applied recent ideas of Smith et al. (2020) and used variation in community composition both within and between ecoregions to determine which areas are most internally homogeneous and most heterogeneous with neighbouring boundaries. We then explored whether internal homogeneity and cross–boundary heterogeneity were driven by their relationships to species replacements and richness differences, and examined how a complex suite of ecogeographical variables (i.e., contemporary environmental features, current climate, topography, Late Quaternary glacial–interglacial climate–change velocity and human footprint) and functional groups (i.e., emergent plants, floating–leaved plants, free–floating plants and submerged plants) affect the robustness of the FEOW ecoregion classification scheme in North American freshwaters.

We hypothesised that **(H1)** ecoregion robustness would decline with increasing latitude (Olson et al., 2001), with ecoregions being least internally homogeneous in their communities (Janzen, 1967), but most dissimilar to one another, in subtropical areas (Sheldon et al., 2018; Smith et al., 2020). Based on a previous study on global variation in community similarity of lake plants (Alahuhta et al., 2017), we also expected **(H2)** that within–ecoregion and across–ecoregion heterogeneity would be caused by species replacements rather than by differences in species richness. Similarly, we predicted **(H3)** that climatic forcing would explain a great deal of variation in the robustness of freshwater plant ecoregions (Heino, 2011; Chappuis et al., 2012; Alahuhta et al., 2021, 2020; García–Girón et al., 2020a, 2020b), with topography, Pleistocene Ice Age legacies, human footprint, water alkalinity, availability of inland waterbodies and the surface area of individual regions playing an important supplementary role (Lacoul & Freedman, 2006; Chappuis et al., 2014; Iversen et al., 2019; Murphy et al., 2019, 2020). Finally, we hypothesised **(H4)** that ecoregions would be a more robust and useful classification for floating–leaved and submerged plants. This derived from the relationships among plant life forms, species–specific tolerance ranges and vagility (Schneider et al., 2018; García–Girón et al., 2019; Gillard et al., 2020), with emergent and free–floating species likely experiencing lower cross–boundary heterogeneity, potentially leading to less defined boundaries in their distributions.

## Materials and Methods

### Ecoregion delineation

The map of freshwater ecoregions that we used for our analyses comes from Abell et al. (2008), which boundaries generally –but not entirely– correspond with those of drainage basins and are roughly equivalent to biomes for terrestrial systems (Abell et al., 2011). The delineation process for North America derives principally from the best available presence/absence information of individual freshwater fish species, coded to eight–digit hydrologic unit codes (HUCs) from NatureServe and published sources. In Canada, separate cluster analyses were conducted on occurrences in each of the nine primary drainage basins, whereas ecoregion delineations in the United States were based on the subregions of Maxwell et al., (1995), with relatively small modifications made following the Endangered Species Committee of the American Fisheries Society (see Abell et al., 2008 for details). As its finest level, this map divides North America (excluding Mexico) into 55 ecoregions and has been shown to be broadly representative of large–scale biogeographical patterns across multiple freshwater organisms, from invertebrates to reptiles (Abell et al., 2008, 2011; Petry et al., 2016). A more detailed description of the delineation methodology is available in Abell et al. (2008) and in their primary sources <https://www.feow.org>.

### Freshwater plant data and explanatory variables

We studied geographical distributions of freshwater vascular plants across North America (from 25°N to 78°N) using a grid of equal–area quadrats, i.e., 50 x 50 km spatial resolution. This dataset is one of the world’s few fine–grained repositories of freshwater plant distributions at continental scales, and has already been described previously to produce maps of species richness (Alahuhta et al., 2020) and investigate range size conservatism and range overlap (Vieira et al., 2021). In brief, distribution maps of 180 freshwater plants were digitalised from the Flora of North America (Flora of North America Editorial Committee, 1993–2007) for a study region that was restricted to the main continental areas of the United States and Canada, excluding Mexico and remote islands. We strictly focused on vascular plant species that are strongly associated with freshwater habitats, removing peatland and marine species following Crow & Hellquist (2000), Flora of North America Editorial Committee (1993–2007), Lichvar (2014) and Murphy et al., (2019, 2020). Hence, riparian, shoreline and semi–aquatic plant species were also excluded from our study. Although we acknowledge that our freshwater species list only consists of a relatively limited number of all aquatic species found in North America (Chambers et al., 2008), all important freshwater hydrophyte genera and species (e.g. *Ceratophyllum* spp., *Myriophyllum* spp., and *Potamogeton* spp.) are present in the data (Crow, 1993, Crow & Hellquist, 2000, Murphy et al., 2019). Moreover, most of the species used in our study have ranges centred in the Northern Hemisphere (Crow, 1993, Chambers et al., 2008), and species richness patterns at continental scales follow those seen at global scales (Murphy et al., 2019, Alahuhta et al., 2020).

We considered ten explanatory variables representative of the ecogeographical variables that we thought most likely would influence the descriptive power of ecoregions in freshwater plants (Alahuhta et al., 2021). These explanatory variables (Supplementary Information Appendix S1) were associated with contemporary environmental features, human footprint, present–day climate, topography, instability of past climate and the surface area of individual regions from the FEOW classification scheme. Here, we used zonal statistics to calculate the mean value for each ecoregion and variable *(sensu* Smith et al., 2020). Environmental features included proportion of freshwaters at 150 m resolution (presence/absence, Lamarché et al., 2017) and mean water alkalinity at 1/16 degrees resolution (mequiv l^−1^, Marcé et al., 2015). Proportion of freshwaters determined the availability of potential habitats for aquatic plants (Jones et al., 2003), whereas alkalinity is a measure of carbon source that can be utilised during photosynthesis (Iversen et al., 2019). Human footprint was assessed based on the global Human Influence Index (HII) from the NASA Socioeconomic Data and Applications Centre <https://earthdata.nasa.gov>. This measure combines metrics of eight variables (i.e., crop land, pasture land, built infrastructure, population density, electric power, roads, railways and navigable waterways) into a single proxy of recent anthropogenic pressures on biodiversity (Sanderson et al., 2002). Current climatic variation indicates not only energy availability and water level fluctuations for freshwater plants, but also material leaching from surrounding lands and potential dispersal events (Kosten et al., 2009; García–Girón et al., 2020a). These variables (i.e., annual mean temperature, °C; annual total precipitation, mm; temperature seasonality, °C; and precipitation seasonality, mmm) were averaged for the period 1970–2000 from WorldClim 2.0 (Fick & Hijmans, 2017), representing both average conditions and their variability across the year. Evidence from recent studies suggests that the extent of mountainous areas is a strong predictor of freshwater plant diversity (Fernández–Aláez et al., 2018) and rarity (García–Girón et al., unpubl). Here, we calculated terrain ruggedness (m) as implemented in the MERIT–Digital Elevation Model (DEM) from the Geomorpho90m global dataset (Amatulli et al., 2020), which uses the NASA Shuttle Radar Topographical Mission (SRTM) to provide topographical variables at 3 arc–second resolution. Although no consensus exists on the influence of Late Quaternary history on freshwater plant diversity (Alahuhta et al., 2020, Murphy et al., 2020), we also calculated the average velocity of climate change from the Last Glacial Maximum (LGM) to present day (expressed as m yr^−1^, i.e., the speed at which species must migrate over the Earth’s surface to maintain constant climatic conditions, Hamann et al., 2015) from a set of transient simulations at *c.* 1/5 degrees resolution (see Sandel et al., 2011 for details).

### Statistical analyses

To determine in what areas ecoregion delineation best describes the underlying variability in freshwater plant distributions (i.e., ecoregion robustness), we calculated **(i)** within–ecoregion homogeneity (i.e., ecoregions that are most internally homogeneous) and **(ii)** cross–boundary heterogeneity (i.e., ecoregions that are highly heterogeneous with nearby areas). First, we used the Sørensen dissimilarity for each individual grid cell, and then averaged the values across the quadrats whose centroids are within the borders of each ecoregion, thereby allowing us to derive a single, ecoregion–level, homogeneity score. Second, we calculated the Sørensen index between all pairs of ecoregions from a presence/absence community matrix synthesising variation in community composition of freshwater plants across North America. Because our analyses here focus principally on the heterogeneity among nearby ecoregions, we followed Smith et al., (2020) and subset these pairwise comparisons to include only comparisons among ecoregions that were fewer than 2,000 km apart. However, since metrics of community distinctness fail to unambiguously capture spatial turnover when they are compared across samples with different species richness (Atmar & Patterson, 1993), we also partitioned the Sørensen index of dissimilarity into its additive fractions (i.e., species replacements and species loss; Carvalho et al., 2012), and checked whether internal homogeneity and across–ecoregion heterogeneity were driven by their relationships to species replacements and richness differences. Importantly, we further stratified our analyses by plant life forms (i.e., emergent plants, floating–leaved plants, free–floating plants and submerged plants; Sculthorpe, 1967; Cook, 1999) to test whether the descriptive power of the FEOW ecoregion classification scheme differed by functional groups (Supplementary Information Appendix S2). We chose to focus on plant life forms not only because information was available for all species (Crow & Hellquist, 2000; Murphy et al., 2019; García–Girón et al., 2020a), but also because these functional categories show differences in their dispersal biology (Santamaría, 2002; García–Girón et al., 2019), as well as in their sensitivity to present–day climate and accessibility to carbon and nutrients from the atmosphere, water and sediments (Lacoul & Freedman, 2006; Alahuhta et al., 2018).

We used spatially explicit regression techniques to examine which explanatory variables contributed most to community dissimilarity within regions and across ecoregion borders. In order to obtain model convergence, we trimmed the original number of candidate variables using multivariate linear regressions. More specifically, we applied forward selection with adjusted R^2^ values (adj. R^2^) and two stopping criteria (i.e., significant level α and global adj. R^2^; Blanchet et al., 2008) to choose statistically explanatory variables to the models (Borcard et al., 2018). Prior to forward selection, we evaluated statistical dependence among the explanatory variables using bivariate correlations (*r* ≥ 0.7; Dormann et al., 2013), transformed these predictors and our response variables to get normally distributed residuals (Peterson & Cavanaugh, 2019), and converted the explanatory variables to their corresponding *z*–scores to allow comparison of their slope coefficients. Both linear and quadratic terms were used in the analyses, not least because we reasonably expected nonlinear impacts of certain explanatory variables. Since Moran’s *I* coefficients using Bonferroni correction (Cabin & Mitchell, 2000) found the presence of spatial autocorrelation in the residuals of preliminary multivariate linear regressions (Supplementary Information Appendices S3 and S4), we constructed simultaneous autoregressive spatial (SAR) models (Cressie, 1993; Haining, 2003). Here, we tested the performance of three different simultaneous autoregressive model types (spatial error model SAR_err_, lagged model SAR_lag_, and mixed model SAR_mix_) and twenty different neighbourhood structures (lag distances between 500 and 10,000 km) with three model selection criteria: **(i)** minimum residual autocorrelation (minimum absolute Moran’s *I* coefficients), **(ii)** maximum model fit (maximum Nagelkerke’s pseudo R^2^_a_), and **(iii)** the Akaike Information Criterion (AIC; Kissling & Carl, 2008).

All statistical analyses were performed in R version 3.6.0 (R Development Core Team, 2018). The list of R packages and statistical routines that have been run throughout this paper is provided in Supplementary Information Appendix S5.

## Results

We found a clear geographical patterning of within–ecoregion homogeneity and cross–boundary heterogeneity in North American freshwater ecoregions, and these results were relatively consistent irrespective of plant life forms (Figures 1 and 2). More specifically, ecoregions were found to be more internally homogeneous and more similar to one another in their communities in polar and subtropical freshwaters, including the northernmost areas of the Canadian Shield, the Arctic Archipelago and the Neotropical Floristic Province of the United States. On the other hand, our models predicted across–ecoregion dissimilarity to be strongest in the temperate floodplain and upland freshwaters in and around the Interior Plains, the Great Lakes and Saint Lawrence region, and the Mediterranean chaparral and endorheic basins of the Southwest, extending along the Pacific Temperate Rainforest and the eastern and western flanks of the Rocky Mountains. Interestingly, the degree of internal homogeneity and ecoregion distinctness were almost equally driven by species replacements (0.26 and 0.32, respectively) and richness differences (0.29 and 0.32, respectively). The replacement component contributed most to ecoregion dissimilarity in the Arctic tundra biome, and species gains and losses more strongly differentiated ecoregions in and around the Great Plain Grasslands (Figures 1 and 2).

**Figure 1.**
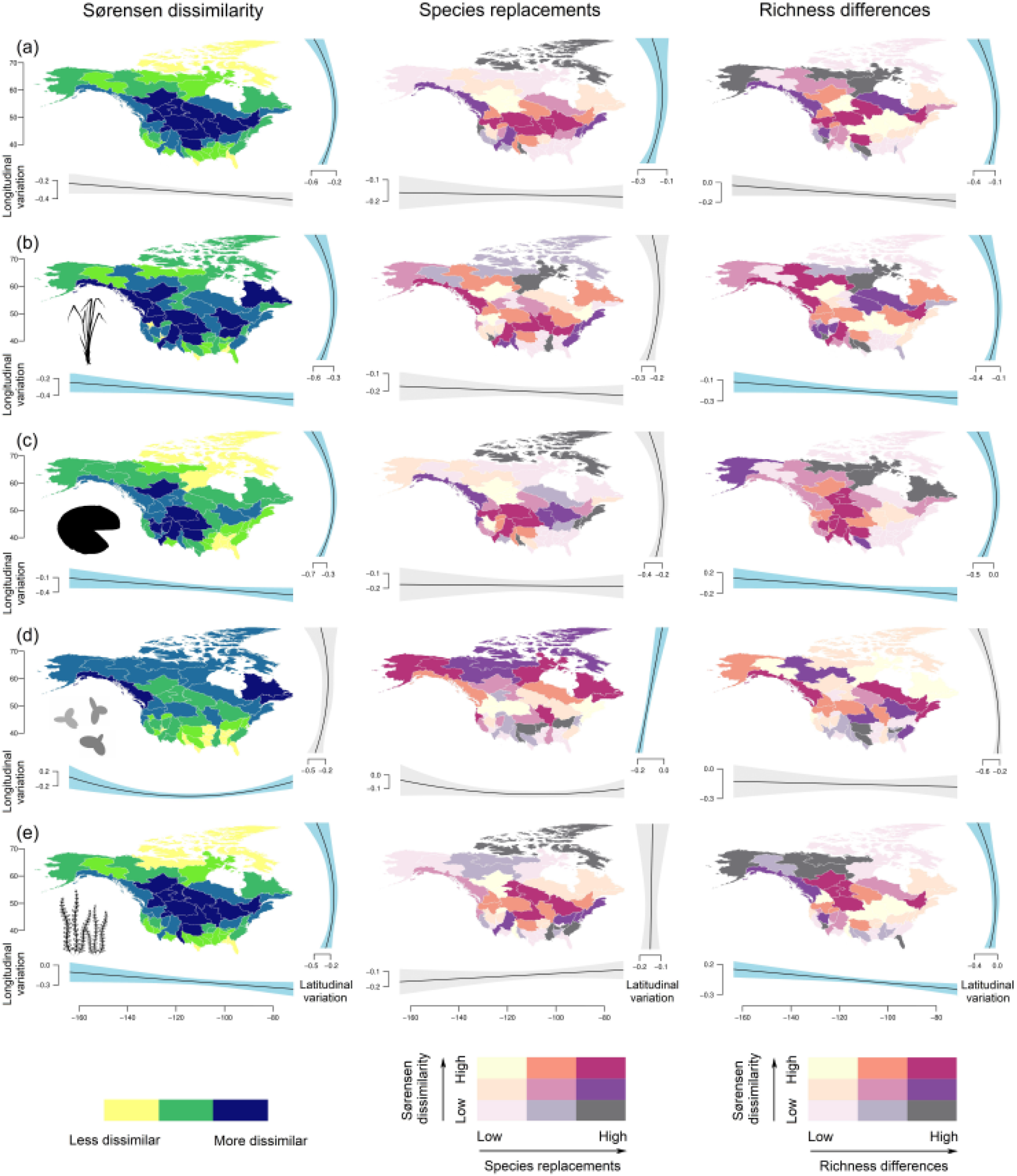
The geographical distribution of within–ecoregion homogeneity for freshwater plant ecoregions in North America (left panels, i.e., darker colours indicate freshwater ecoregions that are less internally homogenous), and bivariate maps comparing internal homogeneity with species replacements and richness differences (centre and right panels, respectively). These maps represent the normalised scores (i.e., the lower the score, the higher the internal homogeneity of individual ecoregions) for **(a)** all freshwater plant species, as well as for analyses stratified by plant life forms, i.e., **(b)** emergent, **(c)** floating–leaved, **(d)** free–floating and **(e)** submerged plants. Relationships of response variables with latitude and longitude were evaluated based on linear regressions, accompanied by Akaike Information Criterion (AIC) to assess the importance of linear vs. quadratic relationships. Solid lines in these plots show the median estimate along with the 95% credible intervals (shading; coloured in blue for statistically significant trends, i.e., *p* ≤ 0.05).

**Figure 2.**
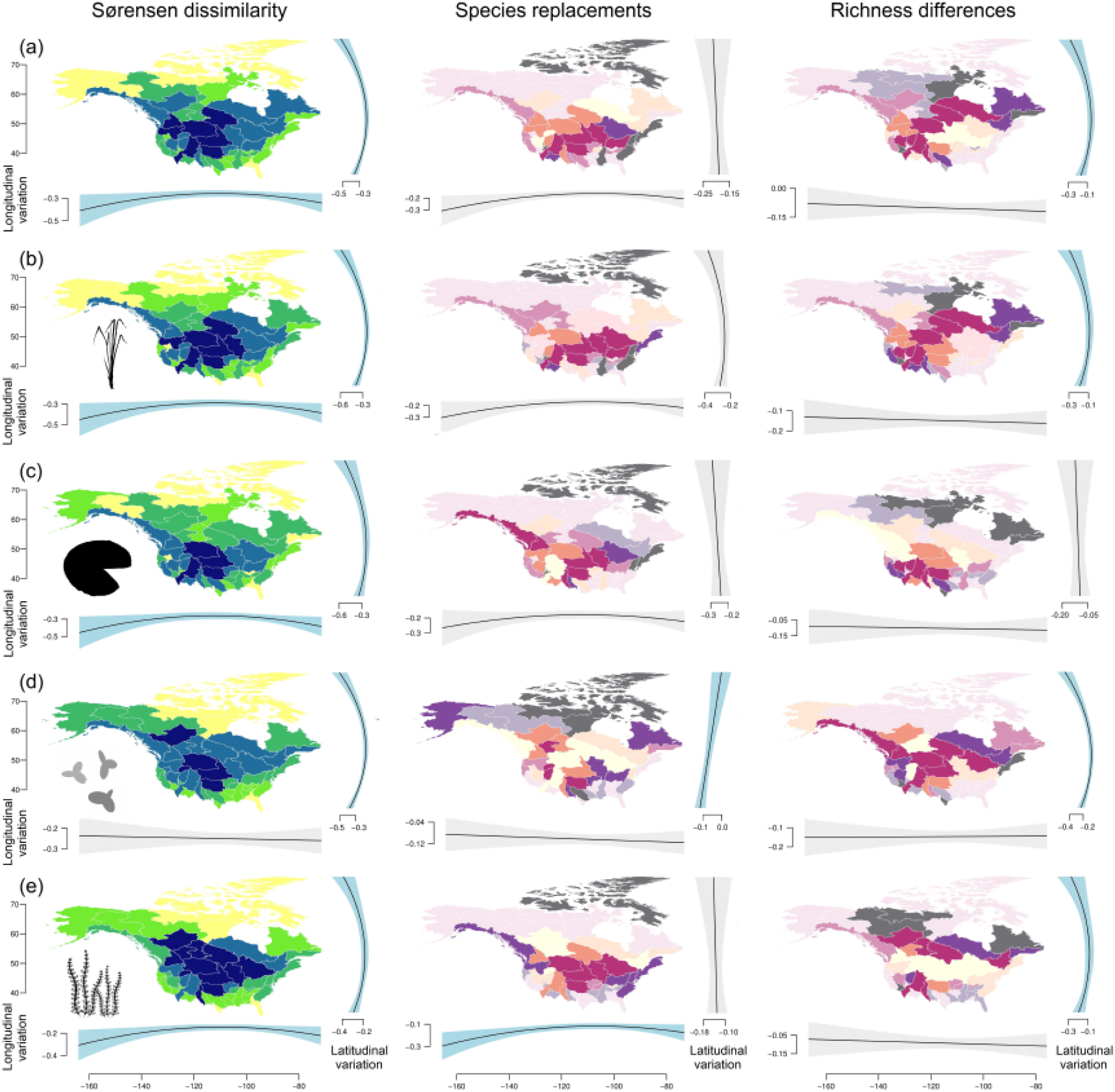
The geographical distribution of cross–boundary heterogeneity for freshwater plant ecoregions in North America (left panels, i.e., darker colours indicate freshwater ecoregions that are more different from one another), and bivariate maps comparing across–ecoregion heterogeneity with species replacements and richness differences (centre and right panels, respectively). These maps represent the normalised scores (i.e., the higher the score, the more dissimilarity exists among ecoregion boundaries) for **(a)** all freshwater plant species, as well as for analyses stratified by plant life forms, i.e., **(b)** emergent, **(c)** floating–leaved, **(d)** free–floating and **(e)** submerged plants. Relationships of response variables with latitude and longitude were evaluated based on linear regressions, accompanied by AIC to assess the importance of linear vs. quadratic relationships. Solid lines in these plots show the median estimate along with the 95% credible intervals (shading: coloured in blue for statistically significant trends, i.e., *p* ≤ 0.05). See Figure 1 for colour scales.

Similarly, we found a strong positive and relatively uniform relationship between the robustness of the FEOW classification scheme and latitude until most northern ecoregions, with ecoregions being less internally homogeneous and more dissimilar to one another at temperate latitudinal bands. Conversely, the relative importance of longitude differed depending on the response variable, with ecoregions being more homogeneous in the East Coast (Figure 1) and cross–boundary heterogeneity being highest in and around the Intermountain region. Not surprisingly, ecoregions predicted to be dissimilar from one another for one functional plant group were also likely to be dissimilar for the other life forms. However, once accounting for the tendency of each life form to have different homogeneity values, ecoregion delineation was more robust for emergent and floating–leaved plants than for submerged hydrophytes, whereas free–floating species tended to be more shared across neighbouring boundaries (Figure 3). In other words, ecoregions were more distinct for emergent and floating–leaved plants than they were for submerged and free–floating hydrophytes.

**Figure 3.**
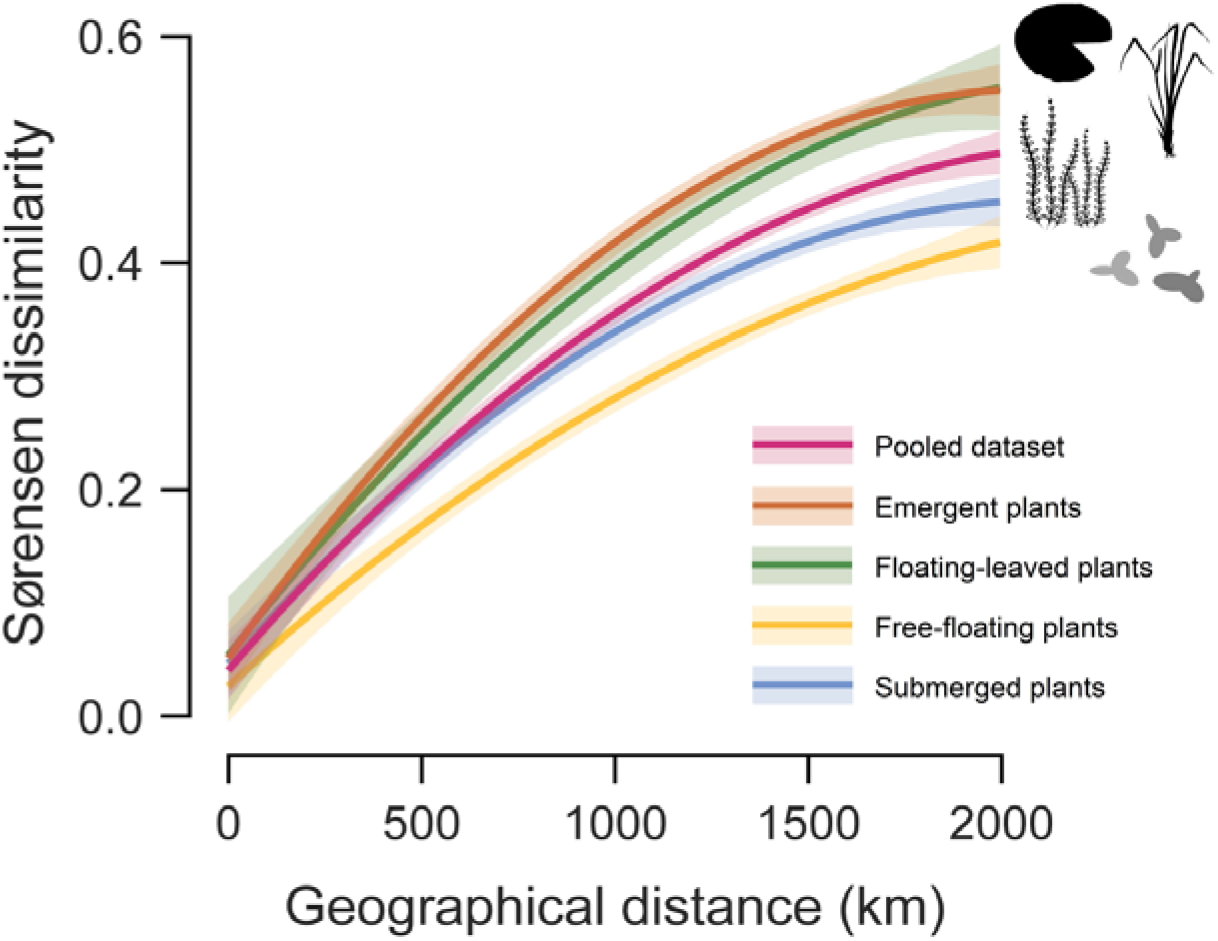
Ecoregion distinctness in relation to geographical distances to neighbouring boundaries (2,000 km; *sensu* Smith et al., 2020). Lines are coloured by plant life form and show the median estimate in a solid line along with the 95% credible intervals (shading), with lower values indicating ecoregions that are more similar in their communities.

After forward selection of explanatory variables (Supplementary Information Appendix S3) in multivariate linear regressions (Supplementary Information Appendix S4), SAR models worked reasonably well, with Nagelkerke’s pseudo R^2^_a_ ranging between 0.10 and 0.74 (Table 1). Despite there was considerable variability in how freshwater plants responded to each individual predictor, some general trends emerged (Figures 4 and 5). Specifically, we found that within–ecoregion homogeneity was highest in regions with higher average temperatures, lower terrain ruggedness, and higher mean water alkalinity concentrations (Figure 4). However, the relative importance of these explanatory variables differed depending on the plant life form considered, with ecoregion area being the best predictor for emergent (Fisher’s Z = 0.47) and free–floating plants (Fisher’s Z = −0.34), and present–day climate being most important for floating–leaved (Fisher’s Z = −0.21) and submerged plants (Fisher’s Z = −0.25). When we examined which variables contributed most to ecoregion distinctness, we found that the three top predictors were the same, i.e., annual mean temperature (Fisher’s Z = −0.42 to −0.53) along with topography (Fisher’s Z = 0.34 to 0.71) and the surface area of individual regions (Fisher’s Z = 0.54 to 0.62; Table 1), with neighbouring boundaries being the most similar to one another at higher annual mean temperatures. Interestingly, there was a strong and positive relationship between increasing terrain ruggedness and increasingly distinct ecoregions. The most surprising relationships involved Late Quaternary Ice Age legacies, with ecoregions more strongly differentiating emergent plant communities in areas that experienced relatively high velocities of climate change since the LGM. Finally, the surface area of individual regions strongly predicted how well the FEOW classification scheme can describe freshwater plant distributions across North America, with medium–sized ecoregions being the most distinct (Figure 5).

**Figure 4.**
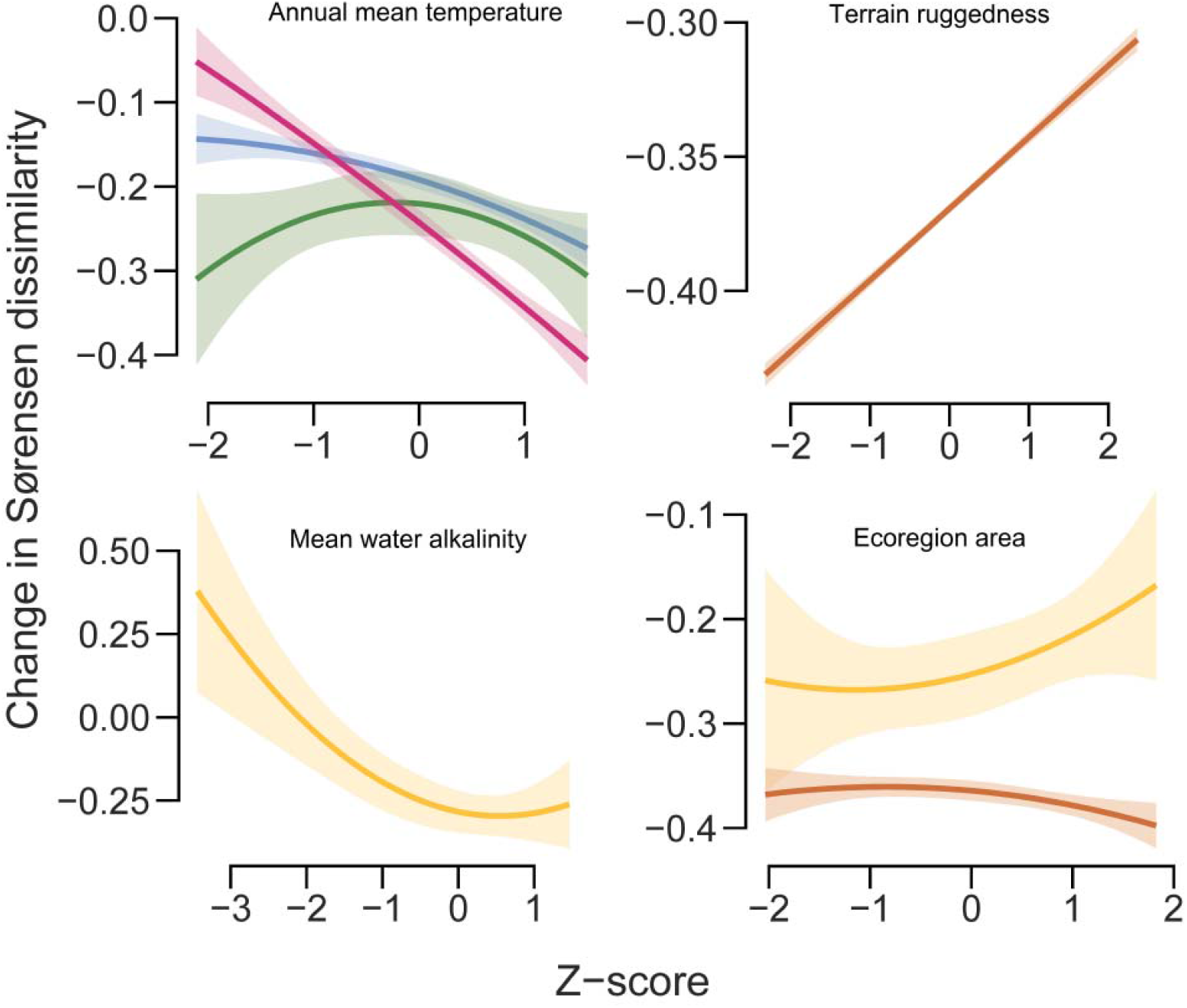
Normalised explanatory variables most strongly related to the internal homogeneity of individual ecoregions. This graph shows the effect of a predictor variable (x–axis) on the changes in the Sørensen dissimilarity index (y–axis). Shown are significant explanatory variables based on forward selection (Supplementary Information Appendix S3) in multivariate linear regressions (Supplementary Information Appendix S4) and spatially explicit regression models (Table 1). Solid lines show the median estimate along with the 95% credible intervals (shading). Ecoregions are most internally homogeneous (i.e., more negative values) in small–sized to medium–sized, warm and flat areas, with their freshwaters having relatively high mean water alkalinity concentrations. See Figure 3 for colour scales of plant functional categories.

**Figure 5.**
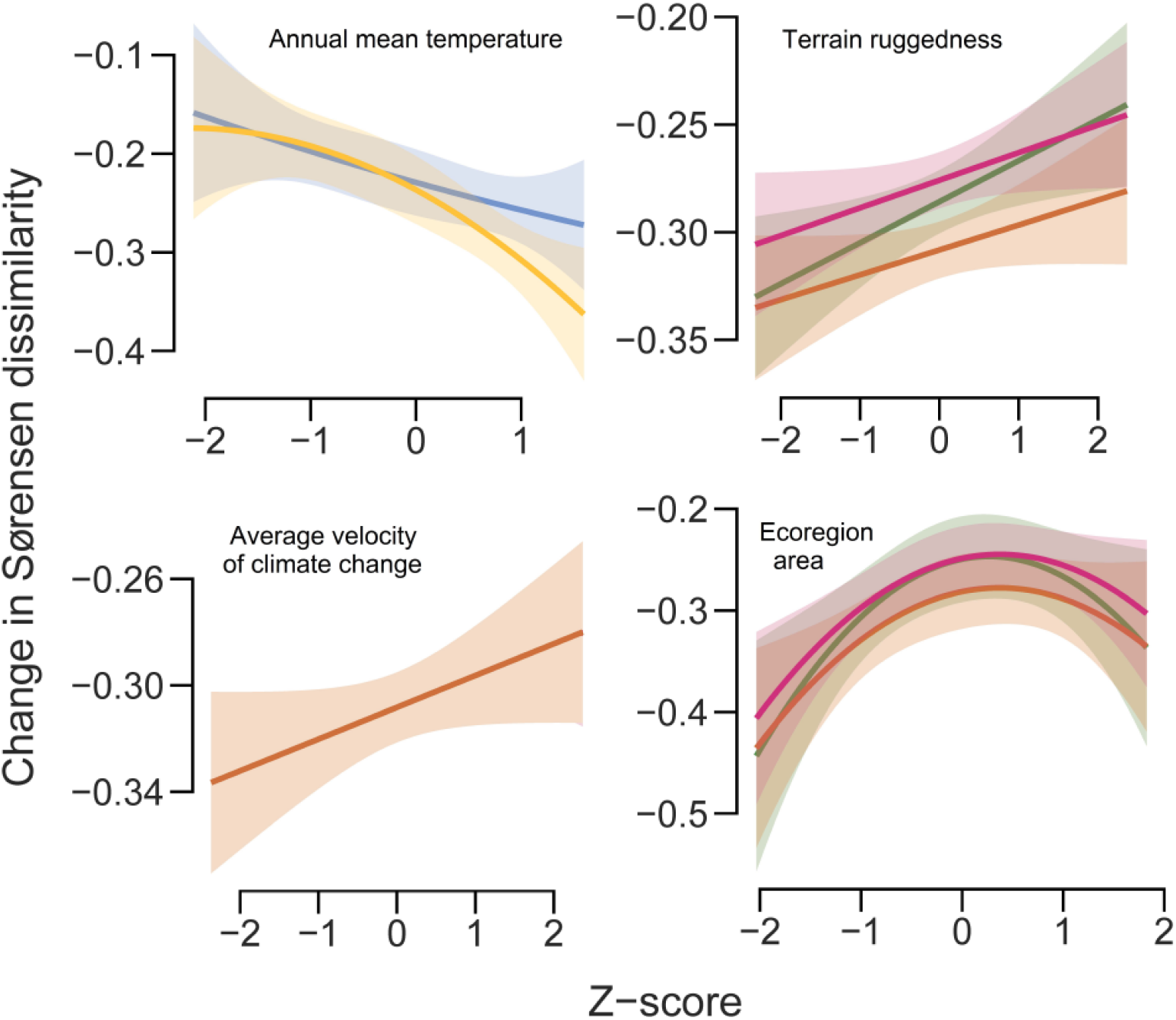
Normalised explanatory variables most strongly related to cross–boundary heterogeneity. This graph shows the effect of a predictor variable (x–axis) on the changes in the Sørensen dissimilarity index (y–axis). Shown are significant explanatory variables based on forward selection (Supplementary Information Appendix S3) in multivariate linear regressions (Supplementary Information Appendix S4) and spatially explicit regression models (Table 1). Solid lines show the median estimate along with the 95% credible intervals (shading). Ecoregions become more distinct (i.e., higher values) in temperate, medium–sized and topographically fragmented landscapes that have experienced oscillations of the Laurentide Ice Sheet after the Pleistocene. See Figure 3 for colour scales of plant functional categories.

**Table 1.**
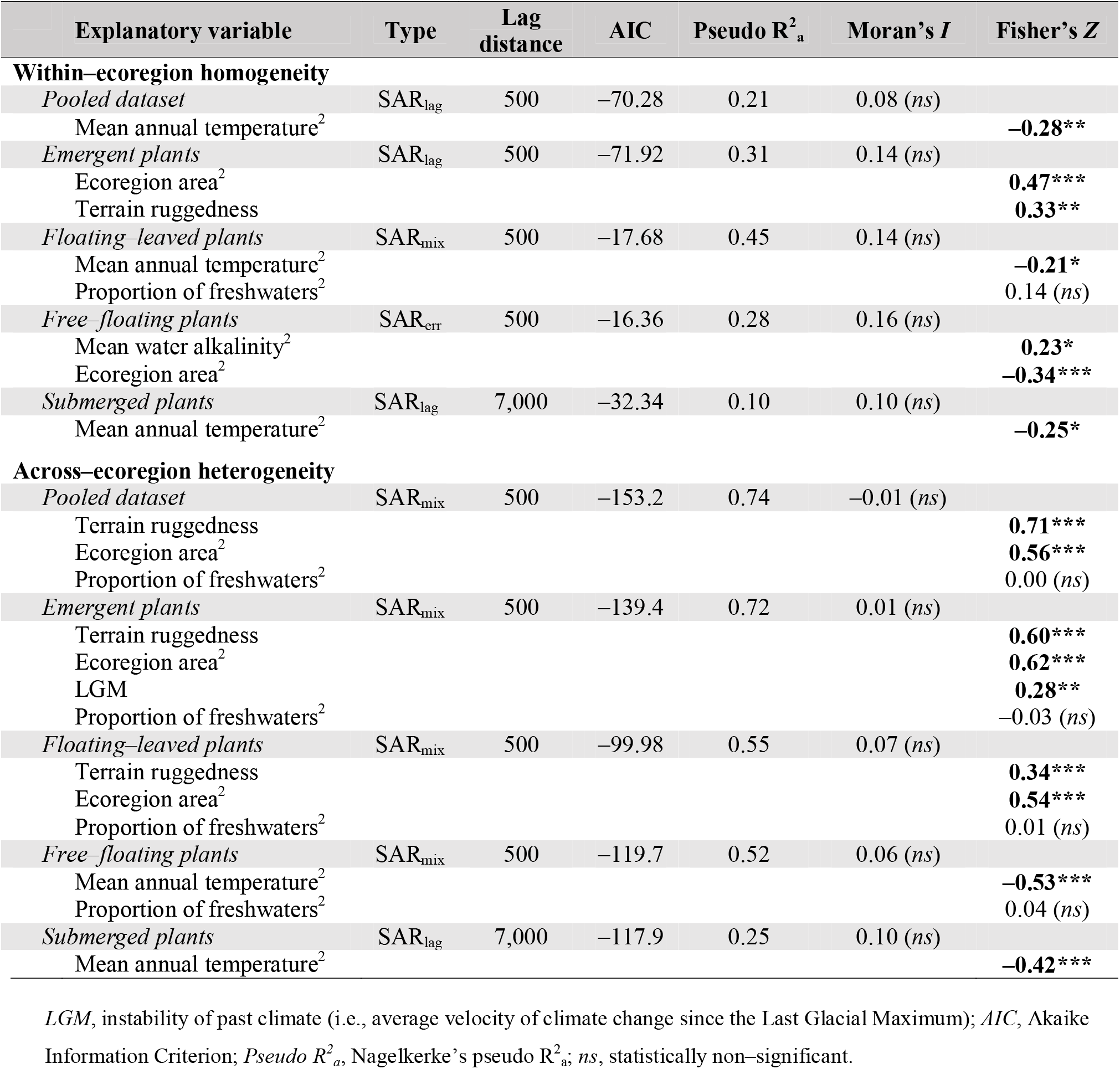
Summary statistics of the ‘best’ simultaneous autoregressive spatial (SAR) models (spatial error model SAR_err_, lagged model SAR_lag_, and mixed model SAR_mix_) explaining internal homogeneity and cross–boundary distinctness of North American freshwater ecoregions. Superscript following an explanatory variable’s name indicates its second order polynomial. Moran’s *I* coefficients for significant lag distances in multivariate linear models (Supplementary Information Appendix S4) were based on Bonferroni correction (Cabin and Mitchell, 2000). The effect size of SAR coefficients (***\*p*** ≤ 0.05; ***\*\*p*** ≤ 0.01; ***\*\*\*p*** < 0.001) was measured using Fisher’s *Z*, which allows for comparison among analyses (Rosenthal, 1994).

## Discussion

Ecoregions can be powerful tools for understanding biodiversity patterns and supporting conservation actions (Dinerstein et al., 2017; Droissart et al., 2018). However, these geographical units are often delineated based on well–known, often charismatic organismal groups, which may not reflect ecoregions for all taxa (Rolls et al., 2017; Ennen et al., 2020). We applied existing freshwater ecoregions of North America founded on fish (Abell et al., 2008) to investigate community dissimilarity of freshwater plants (all taxa and plant life forms separately) and their underlying ecogeographical mechanisms within and across these ecoregions. We based our study on four hypotheses, which received variable degree of support.

Firstly **(H1)**, we assumed that ecoregion robustness declines with increasing latitude (Olson et al., 2001; Sheldon et al., 2018; Smith et al., 2020). We mostly found some support for this hypothesis, although the overall pattern was more hump–shaped than an increasing one, i.e., ecoregions were more internally homogeneous and more similar to one another in their communities in polar and subtropical freshwaters. Secondly **(H2)**, we hypothesised that within–ecoregion and across–ecoregion heterogeneity is caused by species replacements rather than by differences in species richness (Alahuhta et al., 2017). Contrary to our presumption, both mechanisms explained equal amount of variation in internal homogeneity and cross–boundary heterogeneity. Thirdly **(H3)**, we expected climate to have the strongest influence on the variation of freshwater plant ecoregions, followed by other ecogeographical variables (Murphy et al., 2019; García–Girón et al., 2020a). This hypothesis was party supported by our findings, as ecoregion homogeneity and distinctness were best explained by annual mean temperature and terrain ruggedness, respectively, with alkalinity, area and average velocity of climate change having supplementary effects. Fourthly **(H4)**, we hypothesised that ecoregions are a more robust and useful classification scheme for floating–leaved and submerged plants, because emergent and free–floating species likely experience lower across–region heterogeneity (Schneider et al., 2018; García–Girón et al., 2019; Gillard et al., 2020), potentially leading to less defined boundaries in their distributions. For this expectation, we evidenced support for floating–leaved (more robust delineations) and free–floating (less robust delineations) plants, but not for emergent (more robust delineations) and submerged (less robust delineations) taxa.

### Polar and subtropical ecoregions are most internally homogeneous but less distinct from neighbouring boundaries

Most internally homogeneous ecoregions situated in the most southern and northern latitudes. Our outcome partly follows recent evidence gained for a large body of terrestrial taxa, which ecoregions were most homogeneous at tropical and subtropical areas (Smith et al., 2020), although our results also contradict previous findings that ecoregions are more distinct near the Tropics. We speculate that the relatively weak ecoregions distinctness in the southernmost areas of North America originates from their comparatively lower spatial turnover (Figures 1 and 2), an idea that contradicts Janzen’s (1967) original viewpoints on the evolutionary history of tropical and temperate biomes. However, what drives community resemblance in freshwater plants is still largely unknown (Alahuhta et al., 2020), and additional insights from studies conducted in areas outside of North America are needed to provide more empirical foundation for the low across–ecoregion dissimilarity that we found in and around the Subtropics. Interestingly, the found hump–shaped pattern in within–ecoregion homogeneity and cross–boundary heterogeneity closely mimics that of broad–resolution species richness–latitude relationship for freshwater plants in North America (Alahuhta et al., 2020) and worldwide (Murphy et al., 2019). The internal homogeneity of high–latitude ecoregions is speculative though, because the used species data is rather limited in the most northern parts of North America, where vascular plants are presumably substituted by aquatic bryophytes (Heino & Toivonen 2008). Unexpectedly, species replacement and richness difference explained equal amount of variation for ecoregion robustness. Recent studies have clearly shown that species replacement primarily structures freshwater plants independent of spatial scale and study region (Alahuhta et al., 2017; Murphy et al., 2020; Alahuhta et al., 2021). However, these previous exercises utilised an alternative measure of richness difference (i.e., nestedness), which does not consider overall difference in species richness explicitly (Legendre 2014; Schmera et al., 2020). This may partly explain different results between our and other studies.

We further discovered that species replacement and richness difference components varied rather inconsistently even among neighbouring ecoregions with high or low level of ecoregion robustness (Figure 1 and 2). For example, most temperate ecoregions had both high and low degree of species replacement and richness difference in adjacent units. Pinto–Ledezma et al., (2018) detected that species replacement dominated in southern biomes of North America, whereas richness difference (or nestedness in their case) prevailed in the temperate and boreal biomes for (predominantly) terrestrial vascular plants. They further found that species richness was higher in biomes characterised by species replacement and nestedness was more influential in species–poor biomes. These patterns were arguably caused by historical effects and further by differences in speciation time between southern and northern biomes (Pinto–Ledezma et al., 2018). Our findings on freshwater plants give no clear support for their conclusions, as no distinct geographical patterns were generally noticeable for species replacement and richness difference components and species richness, and these two components of beta diversity were not significantly correlated (species replacement: r = −0.02, *p* = 0.91; richness differences: r_s_ = −0.25, *p* = 0.08). Although species replacements outweighed richness differences to some extent in the northernmost ecoregions of Canada, no firm conclusions can be drawn from this due to data limitations at highest latitudes. Historical effects neither had constant influence on ecoregion of freshwater plants in our models. Moreover, Alahuhta et al., (2020) did not report any clear distinction between nestedness and turnover in the range sizes of freshwater plants across North America, but both processes acted simultaneously. This means that, in addition to narrow–ranging species being nested within the distributions of broad–ranging species, some narrow–ranging freshwater plant species are also replaced by broad–ranging species towards high latitudes (Hausdorf and Hennig 2003; Tomasovych et al., 2016).

After analysing multiple organismal groups, Smith et al., (2018) found strong support for the “sharp–transition hypothesis”, where ecoregion borders categorise differentiated biotic communities. Although our analytic approach does not allow us to directly test this hypothesis (or its opposite “gradual transition hypothesis”, where species turnover prevails independent of ecoregion borders; Smith et al., 2018), equal significance of species replacement and richness difference for ecoregion robustness, as well as randomness in the variation of their strength among adjacent ecoregions, implicitly suggests that freshwater plants do not explicitly follow either of the hypothesis. This would further mean that geographical regionalisation based on terrestrial taxa are not applicable for some freshwater biotas. However, more comparative analysis on the validity of sharp–transition vs. gradual transition hypothesis are needed for freshwater plants, not only in North America but also worldwide.

### Climate and topography are the primary drivers of freshwater plant ecoregions in North America

Ecoregions of freshwater plants were mainly driven by current climate (i.e., annual mean temperature) and terrain ruggedness, which had the highest contributions to internal homogeneity and cross–boundary heterogeneity, respectively. Temperature affects physiological responses of freshwater plants (Lacoul and Freedman, 2006), which also suffer from indirect responses of cold temperatures, such as freezing of surface sediments, ice erosion, limiting light penetration and air–water gas exchanges resulting from thick ice and snow cover (Nilsson et al., 2012). Although aquatic ecosystems mitigate extreme atmospheric climate conditions, different present–day climate variables have been evidenced to influence freshwater plant distributions at broad spatial scales (Murphy et al., 2019; Alahuhta et al., 2020; Gillard et al., 2020). On the other hand, topographical variation via terrain ruggedness affected ecoregion robustness of all freshwater taxa, with negative and positive relationships for within–ecoregion homogeneity and across–ecoregion heterogeneity, respectively. Alahuhta et al., (2017) discovered that environmental heterogeneity originated from topographical variation had the highest influence on global beta diversity of lake plants. This observation was further supported by a global study from six continents combining taxonomic, functional and phylogenetic information for lake plant metacommunities (García–Girón et al., 2020a). A greater variety of habitats or resources with greater variation in elevation explained lake plant distributions in their studies, which is consistent with our findings that nearby freshwater ecoregions become more distinct and less internally homogeneous in topographically dissected landscapes.

A recent study resting on over 2 million observations of variable —but mostly terrestrial— biotas reported that ecoregions were most similar to one another at very low or very high levels of human influence, whereas most robust ecoregions were found at intermediate levels of anthropogenic impacts (Smith et al., 2020). We did not detect similar pattern for ecoregion robustness of freshwater plants, as human footprint on natural ecosystems was not selected in any of the models.

### Comparison among plant life forms

Differences in the trends of ecoregion robustness were only modest among different plant life forms, with communities being less internally homogeneous but more distinct to one another at intermediate latitudes. Although these overall geographical patterns were relatively similar among the life forms, we found differences in the strength of ecoregion robustness and ecogeographical variables explaining ecoregion delineations. Floating–leaved and emergent plants showed more robust ecoregions than free–floating and submerged plant taxa, which contradicts our expectation that emergent species show less defined boundaries in their distributions. Emergent and floating plants benefit from a more direct atmospheric connection for carbon and oxygen use (e.g., Iversen et al., 2019), and greater light availability (e.g., Hautier et al., 2009). However, climate or alkalinity did not consistently structure ecoregions of different plant life forms, with the exception of submerged and free floating species, which delineations in terms of internal homogeneity were most robust at higher mean annual temperatures and mean water alkalinity, respectively. Instead, topography and ecoregion area contributed strongest to the ecoregions of floating–leaved and emergent plants. These two variables reflect habitat and ecosystem availability for different species of floating–leaved and emergent plants (Garcia–Giron et al., 2020; Jones et al., 2003). Topographical variation can also indicate a wider range in temperature and precipitation that would support our climate–driven reasoning for these plant groups.

Ecoregions of free–floating plants were more robust at high alkalinities, which was expectable considering that these plants primarily uptake carbon in the form of bicarbonate from water (Iversen et al., 2019). Many submerged species similarly depend on bicarbonate for carbon use, but alkalinity had no significant contribution to them. It may be that stronger effect of climate overshadowed water quality contributions on submerged plants. Our findings also suggested that historical effects contributed to across–region heterogeneity of emergent plants, implying that most diverse ecoregions experience relatively high velocities of climate change since the LGM. However, no similar trend was observed for within–ecoregion homogeneity of emergent taxa, and many of the ecoregions covered by the Laurentide Ice Sheet showed more similarity to one another than more disparate ice–free ecoregions.

In conclusion, our investigation emphasises that geographical regionalisations, such as ecoregions, founded on a particular organismal group are not applicable for all taxa. Ecoregions originally developed for fish did not show consistent robustness for freshwater plants in North America. We found that within–ecoregion homogeneity was highest in regions with higher average temperatures, lower terrain ruggedness, and higher mean water alkalinity concentrations, whereas neighbouring boundaries became more distinct in medium–sized, temperate and topographically fragmented landscapes that have experienced oscillations of the Laurentide Ice Sheet after the LGM. Both species replacement and richness difference components were equally important for ecoregions of freshwater plants but showed no evident geographical trends across the continent. Our study sets first steps for further assessment and development of geographical regionalisations not only for freshwater plants but also for other taxa inhabiting inland water systems. These updated regionalisations can then be used for conserving different biotas in freshwaters, which are currently among the most threatened ecosystems in the world.

## Acknowledgements

Both authors are grateful for the support from Academy of Finland (grants: 322652 and 331957)

## Data availability

Data used in the study will be deposited to Dryad if the manuscript is accepted for publication.

## Biosketch

Janne Alahuhta heads Macrophyte Biogeography Lab operating at University of Oulu and Jorge Garcia□Giron is a postdoc researcher at the Finnish Environment Institute. Both are passionate about biogeography of freshwater assemblages and have actively collaborated to advance our understanding on the patterns and processes underlying broad□scale freshwater biotas since the IBS Malaga 2019 symposium.

JA conceived the original idea, whereas JA and JGG contributed to the study design equally. JGG processed the data and performed the analysis. JA and JGG were together responsible for the writing of the manuscript.

## References

Abell, R., et al. 2008. Freshwater Ecoregions of the World: A New Map of Biogeographic Units for Freshwater Biodiversity Conservation. BioScience, 58, 403–414.

Abell, R., et al. 2011. Concordance of freshwater and terrestrial biodiversity. Conservation Letters, 4, 127–136.

Alahuhta, J., Lindholm, M., Baastrup–Spohr, L., García–Girón, J., Toivanen, M., Heino, J., & Murphy K. J. 2021. Macroecology of macrophytes in the freshwater realm: patterns, mechanisms and implications. Aquatic Botany, 168, 103325.

Alahuhta, J., Antikainen, H., Hjort, J., Helm, A., & Heino, J. 2020. Current climate overrides historical effects on species richness and range size of freshwater plants in Europe and North America. Journal of Ecology, 108, 1262–1275.

Alahuhta, J., et al. 2017. Global variation in the beta diversity of lake macrophytes is driven by environmental heterogeneity rather than latitude. Journal of Biogeography, 44, 1758–1769.

Alahuhta, J., et al. 2018. Global patterns in the metacommunity structuring of lake macrophytes: regional variations and driving factors. Oecologia, 188, 1167–1182.

Amatulli, G., McInerney, D., Sethi, T., Strobl, P., & Domisch, S. 2020. Geomorpho90m, empirical evaluation and accuracy assessment of global high–resolution geomorphometric layers. Scientific Data, 7, 162.

Atmar, W., & Patterson, B. D. 1993. The measure of order and disorder in the distribution of species in fragmented habitat. Oecologia, 96, 373–382.

Blanchet, F. G., Legendre, P., & Borcard, D. 2008. Forward selection of explanatory variables. Ecology, 89, 2623–2632.

Blasi, C., and Frondoni, R. 2011. Modern perspectives for plant sociology; The case of ecological land classification and the ecoregions of Italy. Plant Biosystems, 145, 30□37.

Borcard, D., Gillet, F., & Legendre, P. 2018. Numerical Ecology with R. New York, USA: Springer International Publishing.

Cabin, R. J., & Mitchell, R. J. 2000. To Bonferroni or not to Bonferroni: When and how are the questions. Bulletin of the Ecological Society of America, 81, 246–248.

Carvalho, J. C., Cardoso, P., Borges, P. A. V., Schemera, D., & Podani, J. 2012. Measuring fractions of beta diversity and their relationships to nestedness: a theoretical and empirical comparison of novel approaches. Oikos, 122, 825–834.

Chambers, P., Lacoul, P., Murphy, K. J., & Thomaz, S. M. 2008. Global diversity of aquatic macrophytes in freshwater. Hydrobiologia, 595, 9–26.

Chappuis, E., Ballesteros, E., & Gacia, E. 2012. Distribution and richness of aquatic plants across Europe and Mediterranean countries: patterns, environmental driving factors and comparison with total plant richness. Journal of Vegetation Science, 23, 985–997.

Chappuis, E., Gacia, E., & Ballesteros, E. 2014. Environmental factors explaining the distribution and diversity of vascular aquatic macrophytes in a highly heterogeneous Mediterranean regions. Aquatic Botany, 113, 72–82.

Cook C. D. K. 1999. Aquatic Plant Book (2^nd^ ed.). Amsterdam, NL: SPB Academic Publishing.

Cressie, N. A. C. 1993. Statistics for spatial data. New Jersey, USA: John Wiley & Sons.

Crow, G. E., & Hellquist, C. B. 2000. Aquatic and Wetland Plants of Northeastern North America. Madison, USA: University of Wisconsin Press.

Dias□Loyola, R., Becker, C. G., Kubota, U., Baptista□Haddad, C. F., Fonseca, C. R., Lewinsohn, T. M. 2008. Hung Out to Dry: Choice of Priority Ecoregions for Conserving Threatened Neotropical Anurans Depends on Life□History Traits. PLoS ONE, 3, e2120.

Dormann, C. F., et al. 2013. Collinearity: a review of methods to deal with it and a simulation study evaluating their performance. Ecography, 36, 27–46.

Fernández–Aláez, C., Fernández–Aláez, M., García–Criado, F., & García–Girón, J. 2018. Environmental drivers of aquatic macrophyte assemblages in ponds along an altitudinal gradient. Hydrobiologia, 812, 79–98.

Fick, S. E., & Hijmans, R. J. 2017. WorldClim 2: new 1–km spatial resolution climate surfaces for global land areas. International Journal of Climatology, 37, 4302–4315.

Flora of North America Editorial Committee. 1993–2007. Flora of North America North of Mexico (Vol. 20+). New York, USA: Oxford University Press.

García–Girón, J., et al. 2020a. Global patterns and determinants of lake macrophyte taxonomic, functional and phylogenetic beta diversity. Science of the Total Environment, 723, 138021.

García–Girón, J., et al. 2020b. Elements of lake macrophyte metacommunity structure: Global variation and community–environment relationships. Limnology and Oceanography, 65, 2883–2895.

García–Girón, J., Wilkes, M., Fernández–Aláez, M., & Fernández–Aláez, C. 2019. Processes structuring macrophyte metacommunities in Mediterranean ponds: Combining novel methods to disentangle the role of dispersal limitation, species sorting and spatial scales. Journal of Biogeography, 46, 646–656.

Gillard, M. B., Aroviita, J., & Alahuhta, J. 2020. Same species, same habitat preferences? The distribution of aquatic plants is not explained by the same predictors in lakes and streams. Freshwater Biology, 65, 878–892.

Haining, R. 2003. Spatial data analysis: theory and practice. Cambridge, UK: Cambridge University Press.

Hamann, A., Roberts, D. R., Barber, Q. E., Carroll, C., & Nielsen, S. E. 2015. Velocity of climate change algorithms for guiding conservation and management. Global Change Biology, 21, 997–1004.

Heino, J. 2011. A macroecological perspective of diversity patterns in the freshwater realm. Freshwater Biology, 56, 1703–1722.

Heino, J. & Toivonen, H. 2008. Aquatic plant biodiversity at high latitudes: patterns of richness and rarity in Finnish freshwater macrophytes. Boreal Environment Research, 13, 1□14.

Iversen, L. L., et al. 2019. Catchment properties and the photosynthetic trait composition of freshwater plant communities. Science, 366, 878–881.

Janzen, D. H. 1967. Why Mountain Passes are Higher in the Tropics. The American Naturalist, 101, 233–249.

Jones, J. I., Li, W., & Maberly, S. C. 2003. Area, altitude and aquatic plant diversity. Ecography, 26, 411–420.

Kalwij, J. M., Robertson, M. P., Ronk, A., Zobel, M., & Pärtel, M. 2014. Spatially–explicit estimation of geographical representation in large–scale species distribution datasets. PLoS ONE, 9, e95306.

Kissling, W. D., & Carl, G. 2008. Spatial autocorrelation and the selection of simultaneous autoregressive models. Global Ecology and Biogeography, 17, 59–71.

Kosten, S., et al. 2009. Climate–related differences in the dominance of submerged macrophytes in shallow lakes. Global Change Biology, 15, 2503–2517.

Lacoul, P., & Freedman, B. 2006. Environmental influences on aquatic plants in freshwater ecosystems. Environmental Reviews, 14, 89–136.

Lamarché, C., Santoro M., Bontemps, S., D’Andrimont, R., Radoux, J., Giustarini, L., Brockmann, C., Wevers, J., Defourny, P., & Arino, O. 2017. Compilation and validation of SAR and optical data products for a complete and global map of inland/ocean water tailored to the climate modeling community. Remote Sensing, 9, 36.

Lichvar, R. W. 2014. The National Wetland Plant List: 2014 wetland ratings. Phytoneuron, 41, 1–42.

Marcé, R., Obrador, B., Morguí, J–A., Lluís–Riera, J., López, P., & Armengol, J. 2015. Carbonate weathering as a driver of CO_2_ supersaturation in lakes. Nature Geoscience, 8, 107–111.

Maxwell, J. R., Edwards, C. J., Jensen, M. E., Paustian, S. J., Parrott, H., & Hill, D. M. 1995. A Hierarchical Framework of Aquatic Ecological Units in North America (Nearctic Zone). St. Paul (MN): USDA Forest Service, North Central Forest Experiment Station. General Technical Report NC–176.

Murphy, K. J., Carvalho, P., Efremov, A., Tapia–Grimaldo, J., Molina–Navarro, E., Davidson, T. A., & Thomaz, S. M. 2020. Latitudinal variation in global range–size of aquatic macrophyte species shows evidence for a Rapoport effect. Freshwater Biology, 65, 1622–1640.

Murphy, K., et al. 2019. World distribution, diversity and endemism of aquatic macrophytes. Aquatic Botany, 158, 103127.

Olson, D. M., et al. 2001. Terrestrial ecoregions of the world: A new map of life on Earth. BioScience, 51, 933–938.

Peterson, R. A., & Cavanaugh, J. E. 2019. Ordered quantile normalization: A semiparametric transformation built for the cross–validation era. Journal of Applied Statistics, 47, 2312–2327.

Petry, A. C., et al. 2016. Fish composition and species richness in eastern South American coastal lagoons: additional support for the freshwater ecoregions of the world. Journal of Fish Biology, 89, 280–314.

R Development Core Team. 2018. R: A Language and Environment for Statistical Computing. Vienna, AU: R Foundation for Statistical Computing.

Rosenthal, R. 1994. The Handbook of Research Synthesis. New York, USA: Russel Sage Foundation.

Sandel, B., Arge, L., Dalsgaard, B., Davies, R. G., Gaston, K. J., Sutherland W. J., & Svenning, J–C. 2011. The influence of Late Quaternary climate–change velocity on species endemism. Science, 334, 660–664.

Sanderson, E. W., Jaiteh, M., Levy, M. A., Redford, K. H., Wannebo, A. V., & Woolmer, G. 2002. The human footprint and the last of the wild. Bioscience, 52, 891–904.

Santamaría, L. 2002. Why are most aquatic plants widely distributed? Dispersal, clonal growth and small–scale heterogeneity in a stressful environment. Acta Oecologica, 23, 137–154.

Schneider, B., Cunha, E. R., Marchese, M., & Thomaz, S. M. 2018. Associations between Macrophyte Life Forms and Environmental and Morphometric Factors in a Large Sub–Tropical Floodplain. Frontiers in Plant Sciences, 9, 195.

Sculthorpe C. D. 1967. The Biology of Aquatic Vascular Plants. London, UK: Edward Arnold Publishers Ltd.

Sheldon, K. S., Huey, R. B., Kaspari, M., & Sanders, N. J. 2018. Fifty years of mountain passes: A perspective on Dan Janzen’s classic article. The American Naturalist, 191, 553–565.

Smith, J. R., Hendershot, J. N., Nova, N., & Daily, G. C. 2020. The biogeography of ecoregions: Descriptive power across regions and taxa. Journal of Biogeography, 47, 1413–1426.

Vieira, D. S., García–Girón, J., Heino, J., Toivanen, M., Helm, A., & Alahuhta, J. 2021. Little evidence of range size conservatism in freshwater plants across two continents. Journal of Biogeography, doi: 10.1111/jbi.14071.

